# Quantitative target engagement of RIPK1 in human whole blood via the cellular thermal shift assay for potential pre-clinical and clinical applications

**DOI:** 10.1101/2023.10.02.560523

**Authors:** Shitalben Patel, Marie Karlsson, Joseph T. Klahn, Frank Gambino, Helena Costa, Kathleen A. McGuire, Christina K. Baumgartner, Jon Williams, Sarah Sandoz, James E. Kath

## Abstract

The cellular thermal shift assay (CETSA®) is a target engagement method widely used for preclinical characterization of small molecule compounds. CETSA® has been used for semi-quantitative readouts in whole blood with PBMC isolation, and quantitative, plate-based readouts using cell lines. However, there has been no quantitative evaluation of CETSA® in unprocessed human whole blood, which is preferred for clinical applications. Here we report two separate assay formats – Alpha CETSA® and MSD CETSA® – that require less than 100 μL of whole blood per sample without PBMC isolation. We chose RIPK1 as a proof-of-concept target and, by measuring engagement of seven different inhibitors, demonstrate high assay sensitivity and robustness. These quantitative CETSA® platforms enable possible applications in preclinical pharmacokinetic-pharmacodynamic studies, and direct target engagement with small molecules in clinical trials.

## Introduction

In drug discovery, understanding and quantifying small molecule target engagement is crucial for the characterization of preclinical compounds and clinical drug candidates. Target engagement helps to validate the mechanism of action of lead molecules, determine pharmacokinetic-pharmacodynamic relationships in animal models and trials, and evaluate and mitigate potential off-targets (1,2).

Target engagement can be evaluated by directly measuring small molecule occupancy or by a proximal readout involving pathway changes immediately downstream of the target. Direct and proximal engagement assays are complementary, and both are preferable to a more distal readout that can combine signals from several pathways. While target occupancy assays are routinely used in the development of biologics such as monoclonal antibodies (3), there are fewer options for the evaluation of small molecules, which often target intracellular proteins.

For measuring direct target engagement, the patented cellular thermal shift assay (CETSA®) has become an invaluable tool. CETSA® measures the thermal shift of a target protein when bound to a compound in the relevant cellular environment (4). The underlying principle is that binding of a ligand leads to either a thermal stabilization or destabilization for most proteins, indicating target engagement. The fact that neither the protein of interest nor the test compound needs to be modified with a tag or reporter for CETSA® is a key advantage of the technique. CETSA® can therefore be used in many experimental matrices, such as immortalized and primary cells, tissue sections, and biopsies (5,6). This takes the therapeutically relevant context into account including cell permeability of compound, target protein abundance and localization, and cellular cofactors when measuring target engagement. The resulting data are therefore both physiologically relevant and actionable.

Given these attributes of the CETSA® assay, compound binding can be monitored following *ex vivo* sample treatment or, in animal studies and potentially human trials, following compound dosing. For clinical studies, the most easily accessible sample matrix is blood. There are two proof-of-concept studies that have already described the use of CETSA® with blood, but both utilized peripheral blood mononuclear cell (PBMC) isolation and a semi-quantitative readout (7,8).

RIPK1 (receptor interacting serine/threonine-protein kinase 1) is a key regulator of TNF-mediated apoptosis, necroptosis, and inflammatory pathways that has been associated with autoimmunity as well as inflammatory and neurodegenerative diseases (9,10). RIPK1 inhibitors are currently evaluated in several ongoing clinical trials (11-13). We chose RIPK1 for a proof-of-concept target as its activity in peripheral blood is relevant for disease biology (8), there is a previously described CETSA® assay for use in isolated PBMCs (7), and a variety of reported small molecule inhibitors are commercially available.

Extending on the previous semi-quantitative applications with blood, we developed quantitative CETSA® assays for RIPK1 with two different plate-based readouts. The first technology, AlphaLISA, has previously been combined with CETSA® for assessing direct target engagement in isolated cells (15) and is commonly used in drug discovery; the second, the Meso Scale Discovery electrochemiluminescence assay (MSD), is more commonly used for quantifying biomarkers in clinical trial sample evaluation (*e*.*g*., Refs. 13,14), but to our knowledge has not been used for CETSA®. Our optimized “Alpha CETSA®” and “MSD CETSA®” assays use less than 100 µL of human whole blood per sample and completely circumvent the need for cell isolation steps. The assays correlate well to each other, are robust over several healthy donors, and include a stopping point that is an important prerequisite for multi-site clinical studies.

## Methods

### Compounds

Takeda compound 22, GSK’963, and GSK’962 were synthesized following published protocols (16,17) by WuXi AppTec (Tianjin, China). GSK’963 and GSK’962 are enantiomers and were isolated by preparative chiral supercritical fluid chromatography following synthesis, as described. Necrostatin 4 was additionally synthesized as previously described (18).

The remaining compounds were sourced commercially without additional purification: necrostatin 1 and necrostatin 5 (Cayman Chemical, Ann Arbor, MI, USA), necrostatin 1S (MedChemExpress, Monmouth Junction, NJ, USA), and ponatinib (Advanced ChemBlocks, Hayward, CA, USA). A single lot of each compound was used in all assays; identity and purity (>95% for all) was confirmed for all compounds by LC/MS.

### Statement on Informed Consent

Blood was collected from healthy, anonymized donors into heparin vacutainers or blood bags with citrate phosphate dextrose anticoagulant using venipuncture techniques according to the institutional human subject sample guideline. All subjects provided informed consent.

### Alpha CETSA® with human whole blood

Whole blood, used within one week of a draw, was diluted 1:1 with PBS containing either vehicle control (1% DMSO final concentration) or compound at the indicated concentrations and incubated for 1h at 37 °C. Samples were added to a 96-well PCR plate (10 µL/well) and subjected to heat challenge at temperatures as indicated for 3 minutes using a Veriti Thermal Cycler (Applied Biosystems) with variable temperature zones. After cooling the samples on ice, whole blood samples were further diluted ten-fold with PBS before all samples were lysed by the addition of 5x CETSA® lysis buffer 3, followed by 30 min shaking at room temperature (RT). The assay was performed using a AlphaLISA® SureFire® Ultra RIPK1 kit (Revvity, ALSU-TRIPK1) per instructions.

After sample transfer (5 µL) to a 384-well ProxiPlate (Perkin Elmer, 6008280), 2.5 µL acceptor bead mix (for 300 µL acceptor bead mix: 141 µL RB1, 141 µL RB2, 12 µL activation buffer and 6 µL acceptor beads) was added followed by a 1 h incubation at RT. In the next step, 2.5 µL donor bead mix (for 300 µL donor bead mix; 294 µL dilution buffer and 6 µL donor beads) was added to all wells. AlphaLISA signal was obtained after overnight incubation of detection reagents by measuring the luminescence emission at 615 nm with a PerkinElmer Envision plate reader.

### Preparation of MSD plates for MSD CETSA®

MSD multiarray 96-well plates (Meso Scale Discovery L15XA-3) were coated overnight at 4 °C with 300 ng/mL mouse anti-human RIPK1 capture antibody (1:100 dilution in PBS of BD Bioscience BD610459 lot 0314064, 50 µL/well). Each well was washed three times with 300 μL of wash buffer (PBS pH 7.4, 0.05% Tween-20) and blocked with 150 μL MSD Blocker A (Meso Scale Discovery R93AA-2) for 2 h at RT with shaking at 600 rpm. Plates were finally washed three times with 300 μL wash buffer per well.

### MSD CETSA® with human whole blood

Fresh whole blood was treated with vehicle control (1% DMSO final concentration) or inhibitor at the indicated concentrations. 80 µL per sample was aliquoted into a MicroAmp Optical 96-well PCR plate (ThermoFisher Scientific 4306737) and incubated for 1 h at 37 °C. Blood was heated at temperatures as indicated on a Veriti Thermal Cycler for 3 min. 70 µL per sample was lysed with addition of 10X CST Cell Lysis Buffer (Cell Signaling Technologies 9803S) supplemented with 10X Protease Inhibitor Cocktail (Millipore Sigma P8340) for 30 min at 4 °C while shaking at 600 rpm. Blood can be stored at -80 °C after lysis as a pause point. 60 µL of the lysed blood was further diluted 1:1 with assay diluent (PBS pH 7.4, 1% BSA, 0.05% Tween 20), added to a blocked and washed MSD plate, and incubated overnight at 4 °C with shaking at 600 rpm.

The next day, plates were washed three times with 300 μL wash buffer (PBS pH 7.4, 0.05% Tween-20), 300 ng/mL RIPK1 primary detection antibody (1:300 dilution in Meso Scale Discovery Diluent 100 of Abcam ab125072 lot GR3258748, 50 µL/well) was incubated for 2 h at RT with shaking at 600 rpm, then washed three times with 300 μL of wash buffer. Secondary detection Sulfo-tagged goat anti-rabbit antibody (Meso Scale Discovery, R32AB-1) was diluted 1:1000 in Diluent 100, added to the plate (50 µL/well), incubated for 1 h at RT with shaking at 600 rpm, then washed three times with 300 μL of wash buffer. The assay was developed using 150 uL of MSD read buffer T (R92TC-3, 4X stock diluted 1:1 in water) and read on a MESO Sector S600 plate reader (Meso-Scale Discovery) to measure the electroluminescence signal in each well.

### TEAR1 (Target Engagement Assessment for RIPK1) occupancy assay

RIPK1 occupancy was determined for whole blood incubated 1 h at 37 °C with vehicle control or inhibitor at the indicated concentrations with the orthogonal TEAR1 assay as previously described (14), except the discontinued mouse anti-RIPK1 capture antibody (Abcam ab72139) was replaced with the BD Bioscience 610459 antibody. The normalized unbound RIPK1 ratio for each sample was calculated to determine the % RIPK1 receptor occupancy:

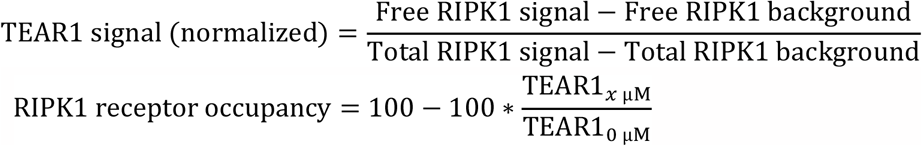

### Data Analysis

Raw signals were plotted and analyzed using GraphPad Prism. CETSA® stabilization values were determined for data at multiple temperatures and one concentration, or a concentration-response at one temperature. For this, the average signal at the maximum temperature condition tested (generally 61 °C for MSD CETSA® and 70 °C for Alpha CETSA®) was subtracted from all assay points, as this was assumed to represent assay background. After background-subtraction, signal at each temperature and concentration point was divided by the average signal at 37 °C vehicle-treated samples. CETSA® signals at a given temperature and inhibitor concentration *x* in some figures were normalized to values at 37 °C and the same inhibitor concentration. These “normalized stabilization” values are therefore corrected for the “non-CETSA® effect” of an inhibitor on the target detection irrespective of heating temperature:

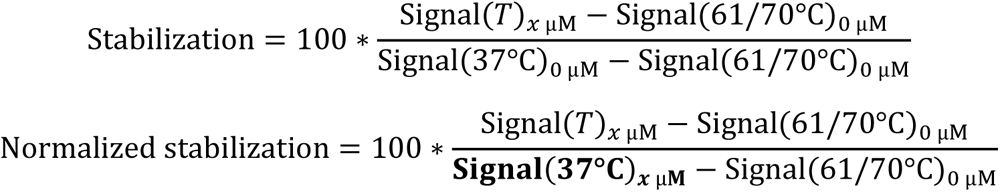

*T*_*m*_, Δ*T*_*m*_, and EC_50_ (melting temperature, difference in melting temperature and half maximal effective concentration) values were calculated from normalized data using nonlinear regression and a 4-parameter fit curve with floating slope, min, and max signal values. Data are presented as mean ± standard error of the mean (SEM). Pearson correlation analysis was performed in GraphPad Prism. The Δ*T*_*m*_ comparisons in Fig. 2 between different assay types, assay runs, and with or without freeze-thaw steps were calculated using unpaired t-test with Welch’s correction. Z’ was calculated from at least 4 replicates of untreated 37 °C heat shocked sample as positive control and untreated 61 °C heat shocked sample as negative control per plate as follows:

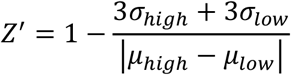

Untreated 37 °C heat shocked and untreated maximally heated (61 °C or 70 °C) samples were used to calculate percent coefficient of variation for inter-donor, intra-donor, and inter-run comparisons.

## RESULTS

CETSA® is commonly performed using semi-quantitative immunoassays, such as immunoblotting or a capillary electrophoresis immunoassay (*e*.*g*., Simple Western), which use a single antibody to quantify the amount of target remaining in a cell’s soluble fraction following a brief heating step and gentle cell lysis. As these immunoassays detect protein under denaturing conditions, the soluble cell fraction must be isolated after lysis in a CETSA® experiment by centrifugation or filtration (Fig. 1A, top schematic).

**Figure 1.**
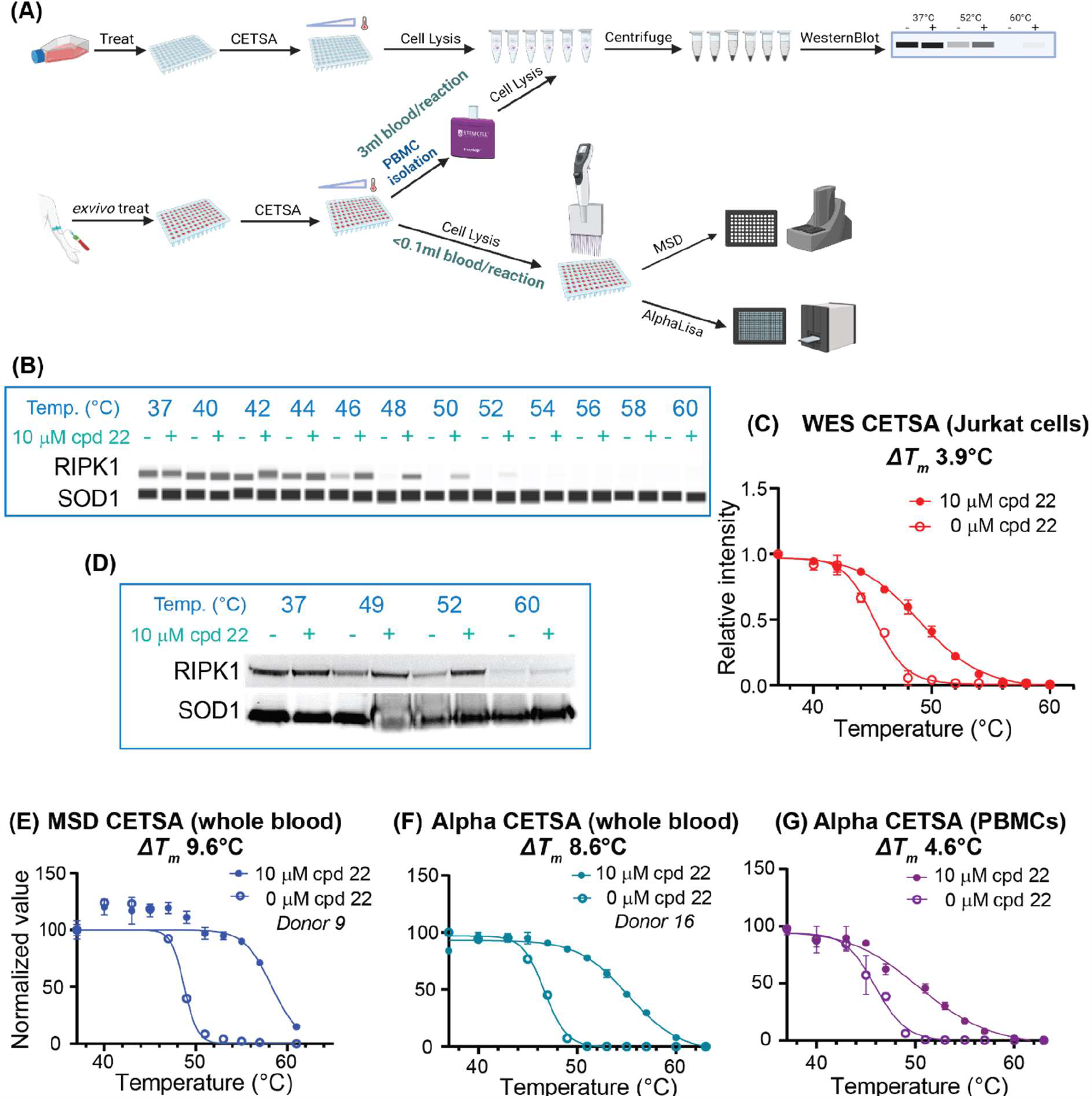
CETSA® formats for cells and whole blood. (A) Schematic for CETSA® with immunoblot (*top*) or AlphaLISA and MSD detection (*bottom*). (B) Simple Western CETSA® signal for soluble RIPK1 in Jurkat cells after treating with 10 μM Takeda compound 22 or vehicle and heating. The thermostable protein SOD1 is also probed as a loading control. (C) Simple Western quantification of RIPK1 CETSA® melting curves in Jurkats. Data are relative to the SOD1 intensity at each temperature and normalized to RIPK1 signal for vehicle at 37°C. (D) Immunoblot CETSA® signal for soluble RIPK1 in whole blood after treating *ex vivo* with 10 μM compound 22, heating, and isolating PBMCs. (E) MSD CETSA® and (F) Alpha CETSA® with whole blood with 10 μM compound 22 and vehicle. (G) Alpha CETSA® data for isolated PBMCs. Data for B, C, D are representative of two biological replicates. 1A was created with Biorender.com.

To establish CETSA® with RIPK1 as a proof-of-concept for assay development, we first treated Jurkat immortalized T lymphocytes with 10 µM Takeda compound 22 (19), a tool inhibitor that has been used for CETSA® experiments with RIPK1 (7). Using the standard CETSA® format with a Simple Western readout, we found that vehicle treated cellular RIPK1 had a typical CETSA® melting curve, with 50% of the protein remaining soluble at a melting temperature (*T*_*m*_) of about 45 °C (Fig. 1B,C). RIPK1 occupied by compound 22, however, had a clear thermal stabilization, with a *T*_*m*_ increase of 3.9 °C (Fig. 1B,C).

A key challenge with human whole blood is that abundant proteins such as albumin, hemoglobin, and immunoglobulins interfere with the above immunoassay formats. In the previously described workflows for CETSA® with blood, this was addressed by isolating PBMCs either before (7) or after (8) the heating step. Using whole blood from a healthy donor, we found PBMC isolation using an EasySep Isolation Kit enabled us to detect a thermal stabilization of RIPK1 in *ex vivo* treated blood (Fig. 1D) in line with a previous report (7). However, this approach, including the use of immunoblotting, remains semi-quantitative and requires about 3 mL whole blood per sample.

For CETSA® to add value to preclinical and clinical studies with whole blood, robust and easily adopted detection methods are needed. Two options we considered were AlphaLISA and MSD. Both have high sensitivity and specificity, a low sample volume requirement, and work well with diverse biological matrices. For either methodology, using an antibody pair that selectively detects protein in a native or soluble state eliminates the need for a centrifugation or filtration step that would be challenging to include in clinical trial sample processing. While this has already been established when applying AlphaLISA for cell-based CETSA® assays (15), it has not been demonstrated in whole blood. Additionally, there is no report of MSD-based CETSA® in any sample type.

For AlphaLISA, we selected a previously established RIPK1 CETSA® assay based on an AlphaLISA® SureFire® Ultra RIPK1 kit from Revvity, previously used for CETSA® with immortalized cell lines (20). As MSD had not been applied to CETSA®, we evaluated a previously described application of the technology to quantify total RIPK1 abundance in human whole blood as part of a separate engagement assay called TEAR1 (Target Engagement Assessment for RIPK1, ref. 14, described below). Assay optimization included antibody dilution (for MSD), lysis buffer selection, temperature, and dynamic range verification. Both assays were optimized first on isolated PBMCs or Jurkat cells (Fig. S1) and validated on RIPK1 wildtype and knockout cell lines. In each case, we confirmed that immunoassay signal was dependent on RIPK1 and was abrogated by heat-induced RIPK1 unfolding or aggregation (Fig. S2).

In healthy human whole blood, we found that the AlphaLISA and MSD CETSA® assays (“Alpha CETSA®” and “MSD CETSA®”) maintained excellent signal/background (S/B) values when comparing maximally folded or denatured RIPK1 (Fig. S3, SI Table 1). Both readouts were in their respective linear ranges for detection (Fig. S3B-E), demonstrating compatibility with this complex biological matrix. In the case of MSD CETSA®, we found that the optimal lysis buffer was different for whole blood compared to cell samples (Fig. S3A). In both cases, we also determined that diluting blood 1:1 with PBS either immediately before (Alpha CETSA®) or immediately after the heat shock (MSD CETSA®) also improved S/B (Fig. S3A and data not shown).

**Table 1.** RIPK1 inhibitors tested with Alpha CETSA®, MSD CETSA®, and the TEAR1 occupancy assay and EC_50_ values.

| **RIPK1 Inhibitor** | **RIPK1 Inhibitor Type** | **Alpha CETSA ITDR 52 °C EC<sub>50</sub>, μM (N, donors)** | **MSD CETSA ITDR 52 °C EC<sub>50</sub>, μM (N, donors)** | **MSD TEAR1 EC<sub>50</sub>, μM (N, donors)** |
| --- | --- | --- | --- | --- |
| Compound 22 | Type III | 0.78 (N=9) | 0.02 (N=2) | 0.028 (N=2) |
| GSK'963 | Type III | 0.48 (N=2) | 0.18 (N=2) | 0.003 (N=2) |
| GSK'962 | Type III | >100 (N=2) | >100 (N=2) | 1.01 (N=2) |
| Necrostatin 1S | Type III | 4.78 (N=2) | 2.51 (N=2) | 0.24 (N=2) |
| Necrostatin 1 | Type III | 22.59 (N=2) | 13.81 (N=2) | 2.13 (N=2) |
| Necrostatin 4 | Type III | 35.95 (N=2) | 48.59 (N=2) | 96.95 (N=2) |
| Necrostatin 5 | Type III | NV | NV | 0.34 (N=1) |
| Ponatinib | Type II | 32.94 (N=2) | 19.85 (N=2) | 29.87 (N=2) |

NV = No Value could be determined from the concentration range tested

For both assays, we observed an absolute reduction in raw RIPK1 detection signal in the presence of compound 22, independent of the heating step – a small reduction for MSD CETSA® (Fig. S3C) and a moderate one for Alpha CETSA® (Fig. S3E). This “non-CETSA® effect” can be observed for immunoassays which detect folded protein and is often attributed to an inhibitor-induced conformational change influencing binding to the target epitope – in extreme cases, an inhibitor can completely block antibody detection (*e*.*g*., in ref. 14, see discussion of the TEAR1 assay below). Although we cannot definitively attribute the non-CETSA® effects for these assays to a RIPK1 conformational change, we still see a similar reduction when adding compound after blood lysis (Fig. S3F), suggesting it is not due to a compound-dependent reduction in RIPK1 abundance. Importantly, the non-CETSA® effect can be corrected by normalizing soluble target signal at each temperature in the presence of compound to the signal at 37 °C, also in the presence of compound (see Methods).

Using the resulting optimized protocols (Fig. 1A, bottom schematic), we generated normalized, temperature dependent CETSA® curves for RIPK1 in heated whole blood following *ex vivo* treatment with 10 μM compound 22 or vehicle control (Fig.1E,F). Despite representing two immunoassays using different detection methods (Alpha CETSA® and MSD CETSA®) and being performed at different labs with different blood donor pools (in Sweden and the United States), we found strikingly similar melting curves and thermal stabilizations for RIPK1 in whole blood (*T*_*m*_ and Δ*T*_*m*_, determined from curve fits, see Methods). Surprisingly, in both cases, the stabilization of RIPK1 after incubation with compound 22 is more pronounced for MSD CETSA® in whole blood (Fig. 1E, Δ*T*_*m*_ 9.6 °C) than in Jurkat cells (Fig. 1B,C, Δ*T*_*m*_ 3.9 °C), and for Alpha CETSA® in whole blood (Fig. 1F, Δ*T*_*m*_ 8.6 °C) than in PBMCs (Fig 1G, Δ*T*_*m*_ 4.6 °C). Although the exact cause of the effect is unclear, thermal stabilization of RIPK1 in whole blood is advantageous to similar measurements with isolated cells.

Blood is a complex biological matrix and the cell composition can vary for different individuals and even for the same individual over time -which may be attributed to conditions like infections or inflammatory diseases. Therefore, both assays were tested on multiple healthy donors, using *ex vivo* treatment with 10 μM compound 22 and heating blood at multiple temperatures to understand the degree of donor-to-donor variability as well as overall assay robustness.

Across five healthy donors for Alpha CETSA®, we found that the inter-donor *T*_*m*_ range within vehicle and compound 22 treated samples was small with values ranging across a temperature span of 0.54 °C and 2.7 °C, respectively. However, the stabilizing effect of compound 22 compared to vehicle control provided a wide assay window with a thermal shift of 8.53 ± 0.37 °C (mean ± standard error [SE]) (Fig. 2A). For MSD CETSA® across four donors, the *T*_*m*_ variability was similar for vehicle and compound-treated samples (value ranging across a temperature span of of 0.32 °C and 1.2 °C, respectively) relative to the stabilization effect (8.60 ± 0.31 °C) (Fig. 2B). The greatest source of variation appears to be present in the compound-treated samples. Overall, this data demonstrates reproducible and robust Δ*T*_*m*_ determination for a tool RIPK1 inhibitor across multiple donors and between the two assays.

**Figure 2.**
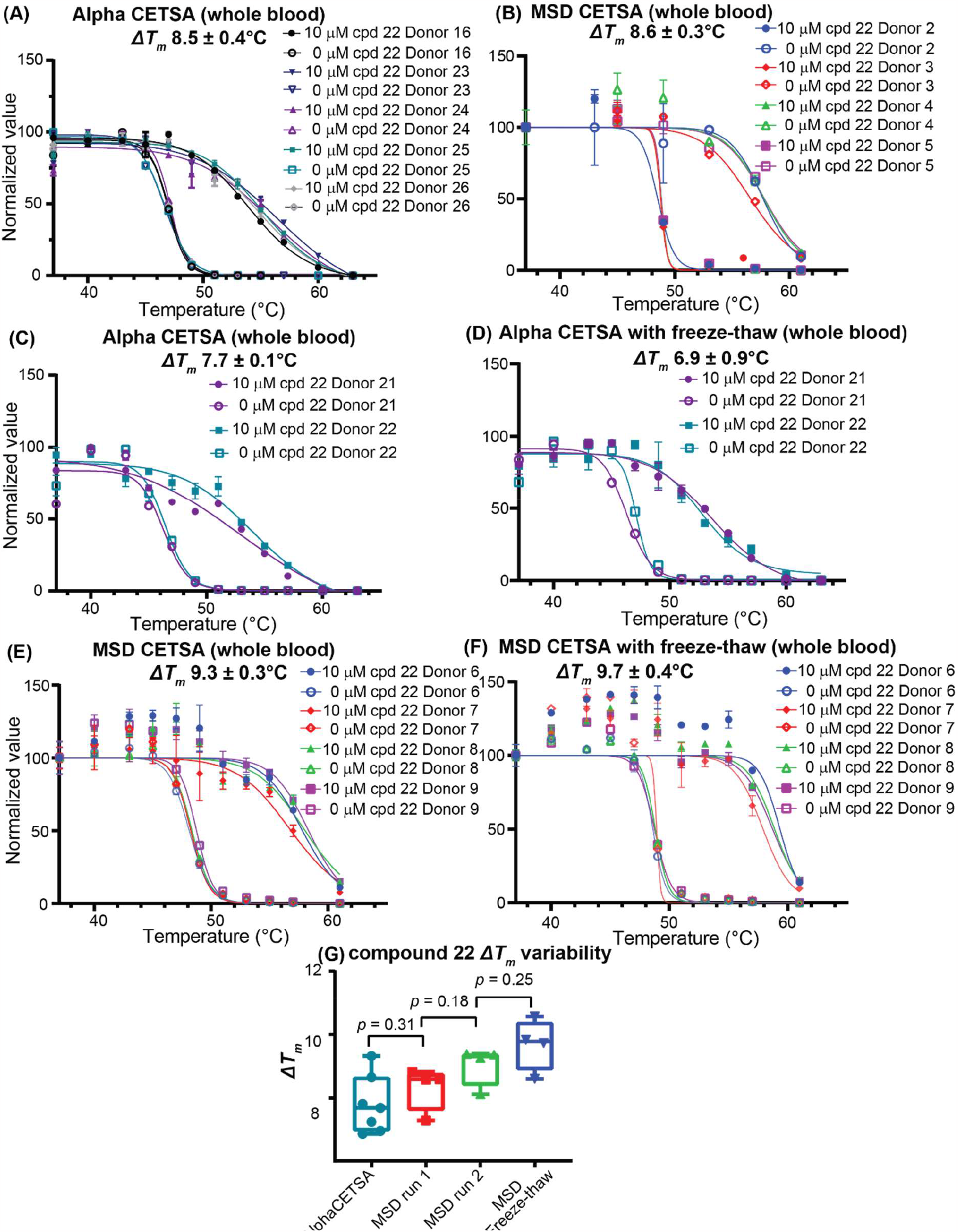
CETSA® thermal shifts (Δ*T*_*m*_) for RIPK1 with Takeda compound 22 in human whole blood from multiple healthy donors. Measurements with (A) Alpha CETSA® or (B) MSD CETSA®. Comparability of CETSA® melting and shift curves with and without a freeze-thaw pause step for (C,D) Alpha CETSA® and (E,F) MSD CETSA®. (G) Comparison of compound 22 Δ*T*_*m*_ variability between Alpha CETSA® (blue) and MSD CETSA® (red) assays across multiple, different donors; between different sets of donors analyzed with MSD CETSA® on different days (red and green); and the same donor sample set analyzed using MSD CETSA® without (green) or with (blue) a freeze-thaw pause step.

To further characterize the assays, we used a calculation of the Z-factor (Z’), a measure of assay performance based on the ratio of the signal standard deviation to the signal window (21). For CETSA®, the signal window can be determined from the highest signal (blood incubated at 37°C) and the lowest signal (blood heated at 61°C or above to fully denature and/or aggregate RIPK1). For Alpha CETSA® performed in Sweden and MSD CETSA® performed in the United States, the Z’ values were 0.76 ± 0.05 (N=6 plates) and 0.78 ± 0.02 (N=33), respectively (SI Table 1 and Fig. S4). Generally, a Z’ above 0.5 is considered acceptable for a quantitative assay, with values approaching 1 representing theoretically ideal assays (21). Additionally, the coefficient of variation, calculated from the mean and SE was less than 10% for different donors between MSD runs calculated from more than 100 high and low signal samples (SI Table 1). The inter-, intra-donor, and day-to-day variability was acceptable and demonstrates that the quantitative Alpha and MSD CETSA® assays for RIPK1 in whole blood are reliable and maintain acceptable technical and biological variability throughout their use.

In order for a direct target engagement assay to be amenable to the clinic, it is invaluable to have a stopping or “pause” point in the workflow. This would allow for asynchronous collection from multiple trial participants and clinical trial sites and enable samples to be processed together at a centralized site equipped with reagents and instruments for MSD or AlphaLISA. We determined that freezing blood after heating and lysis might be an suitable for a pause point since freeze thaw cycles commonly assist the gentle lysis in CETSA® protocols (4) and typically do not affect CETSA® results.

We directly tested this for RIPK1 with human whole blood from two separate donors and found no significant difference between Takeda compound 22 Δ*T*_*m*_ values in either directly processed or frozen samples measured by Alpha CETSA® (Fig. 2C,D) or MSD CETSA® (Fig. 2E,F). Similarly, we did not find any significant difference in the Δ*T*_*m*_ values between the two assays or, for the MSD assay, separate sets of donors processed and analyzed on separate days (Fig. 2G). These data confirm that blood samples for a CETSA® target engagement study can be frozen after the lysis step to pause the assay.

Comparing Δ*T*_*m*_ values between donors is a good measure of the reproducibility and stability of the assay. However, as we see a greater inter-donor range in the *T*_*m*_ for compound-stabilized RIPK1 compared to vehicle-treated samples, we might also expect a significant variation among EC_50_ values in concentration-response (CR) CETSA® experiments determined at a single optimized temperature, also known as isothermal dose response CETSA® (4).

We therefore first treated human whole blood *ex vivo* with concentrations of compound 22 ranging from 100 pM to 100 µM. In MSD CETSA®, we were able to detect CR RIPK1 stabilization with a large window at 53 °C (Fig. 3A). This temperature was then used to determine the CR EC_50_ of compound 22 across four healthy donors (EC_50_ = 0.011 ± 0.001 μM Fig. 3B). For Alpha CETSA®, temperatures from 49-51 °C were tested (Fig. S5) and 50 °C was chosen for EC_50_ determination. The Alpha CETSA® EC_50_ for compound 22 was 0.042 ± 0.011 µM across six healthy donors (Fig. 3C). These data demonstrate a low inter-individual variability for the target engagement measurement across readouts.

**Figure 3.**
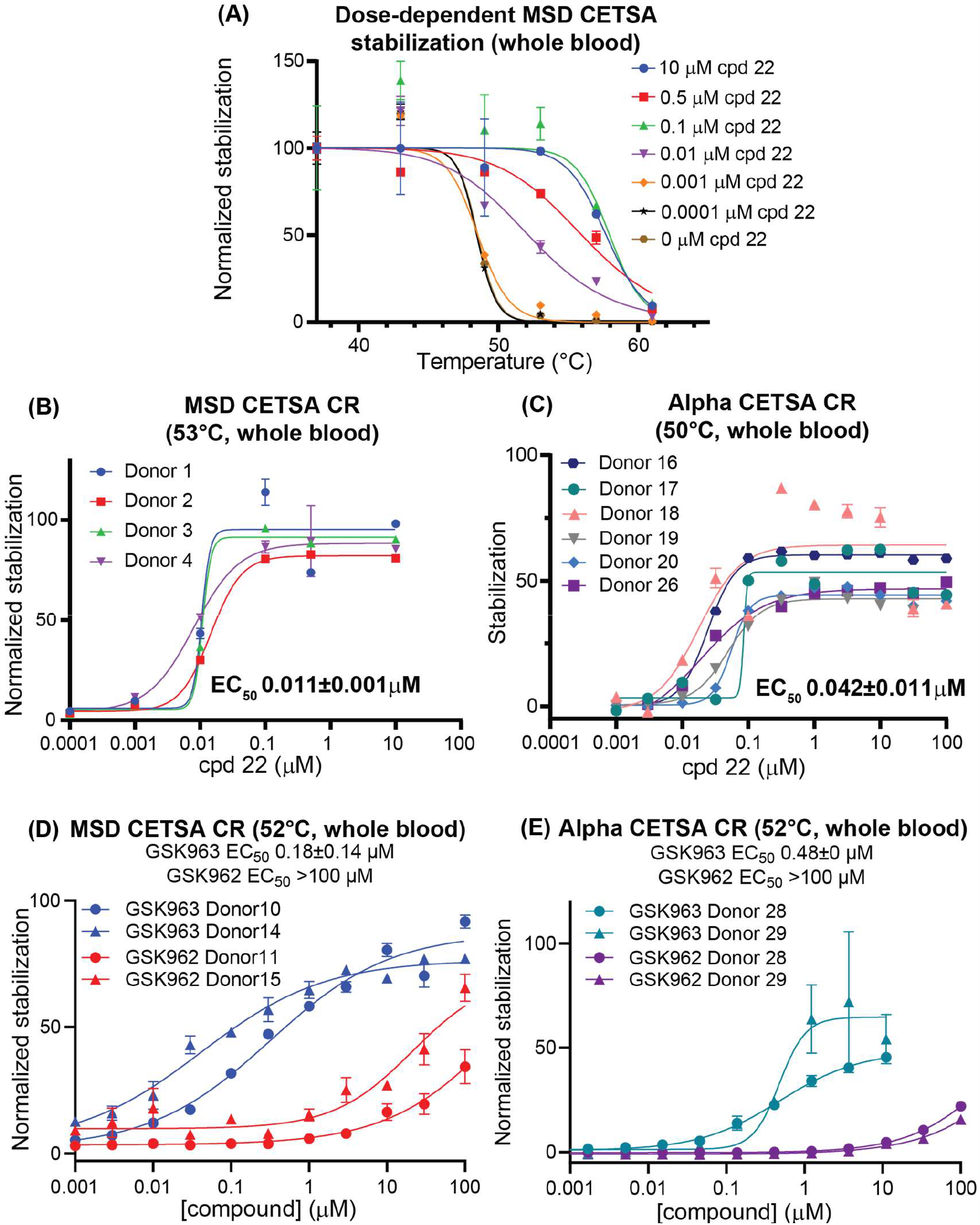
Concentration-response (CR) CETSA® with whole blood. (A) CETSA® melting curves of whole blood with MSD CETSA® at 6 different concentrations of Takeda compound 22 or vehicle. CR CETSA® stabilization of RIPK1 by compound 22 with (B) MSD CETSA® in whole blood from four healthy donors at 53 °C or (C) Alpha CETSA® in whole blood from six donors at 50 °C. Comparison of active and inactive enantiomers GSK’963 and GSK’962 with whole blood from multiple donors with (D) MSD CETSA® or (E) Alpha CETSA®.

For a subset of donors, we determined the reduction of RIPK1 detection signal in the CR readout in the absence of the heating step. As in experiments with a single concentration of compound 22, we saw a minimal effect in the MSD CETSA® assay (Fig. S6A); however, we saw a larger reduction in signal for the Alpha CETSA® format with two additional donors tested (Fig. S6B,C). Despite this larger compound-dependent reduction in detection signal, correcting for the non-CETSA® effect only modestly changed the EC_50_ determined in Alpha CETSA® experiments. When repeating the CR experiments for compound 22 with a different donor pool at a higher temperature, we observed a higher inter-donor variability (Fig. S6D and SI Table 1).

Continuing the evaluation of these quantitative CETSA® assays, we tested additional RIPK1 tool inhibitors with human whole blood while correcting for the non-CETSA® effect. One such compound is GSK’963, along with its inactive enantiomer, GSK’962 (16). By testing GSK’963 at different temperatures and inhibitor concentrations with the MSD CETSA® assay, we found that 52 °C or 53 °C gave similar, large windows for compound-induced thermal stabilization (Fig. S7). In CR experiments, we found, as expected, a large increase in EC_50_ for the inactive enantiomer GSK’962 relative to GSK’963 in Alpha and MSD CETSA® (EC_50_ > 100 μM for GSK’962 in both assays, Fig. 3D,E).

To expand the diversity of RIPK1 inhibitors evaluated, we added necrostatin 1, a well-established tool compound (22), its analogs necrostatin 1S (22), necrostatin 4 (18), and necrostatin 5 (23), as well as ponatinib (24), a less selective inhibitor with activity against RIPK1 (7). In contrast to all the other compounds, which have a unique allosteric type III binding mode, ponatinib is a type II kinase inhibitor. We completed CR experiments with whole blood for these eight compounds at 52 °C (Fig. 3D,E, S8A,B, and Table 1). We were able to determine EC_50_ values for all compounds except necrostatin 5, which did not significantly stabilize RIPK1 even at 100 μM. For the remaining seven compounds, we found a strong correlation for the EC_50_ values between the two assays, with a Pearson coefficient of 0.94 (Fig. 4A, Table 1).

**Figure 4.**
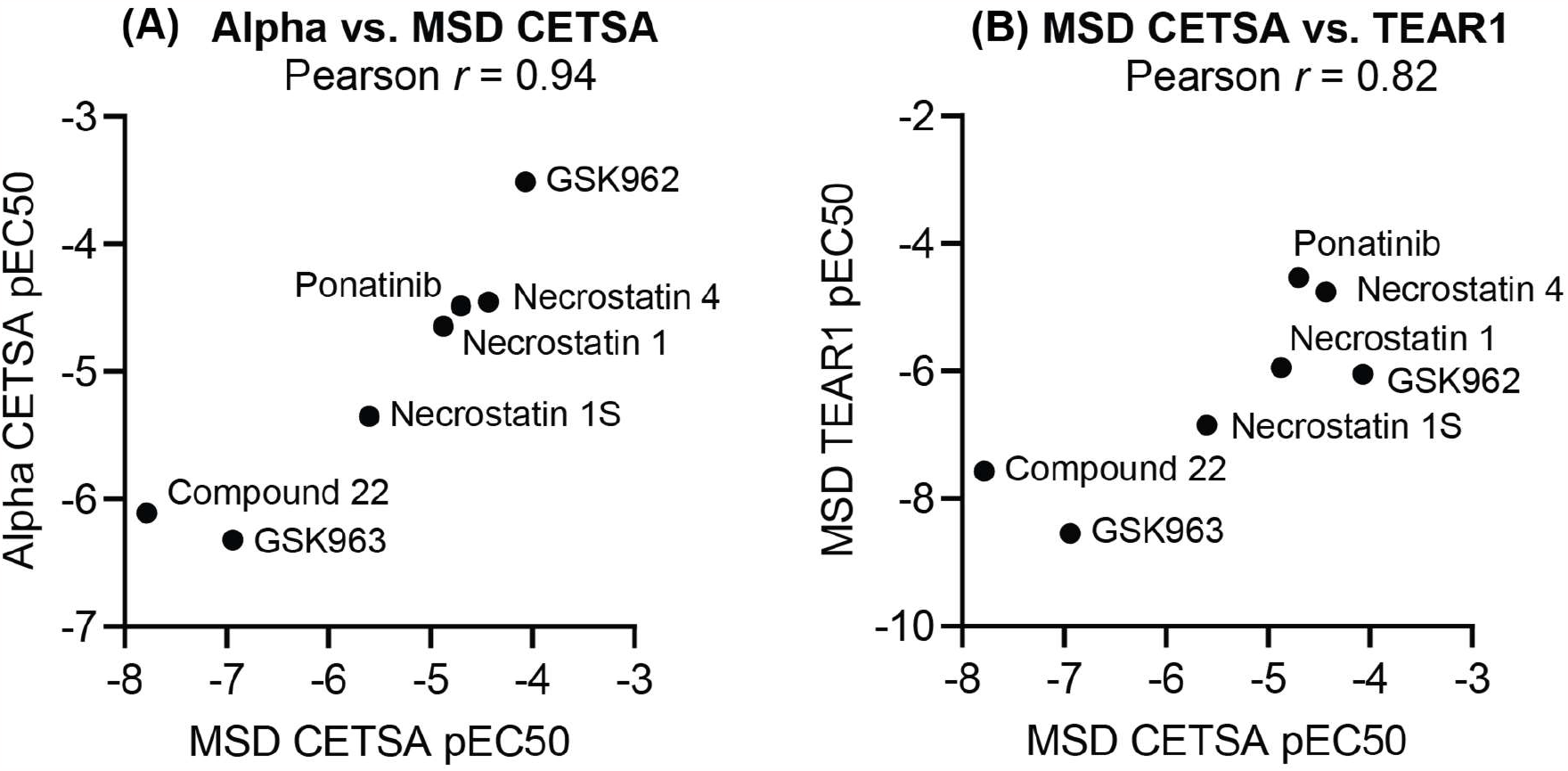
Correlations for engagement measurements across multiple RIPK1 inhibitors. Comparisons for (A) Alpha CETSA® and MSD CETSA® and (B) MSD CETSA® and the TEAR1 assay.

Finally, we decided to compare whole blood CETSA® with the TEAR1 assay, a distinct MSD-based occupancy assay that has been previously developed (14). The TEAR1 assay is similar to a receptor occupancy assay used for biologics, where one antibody pair (“FREE-RIPK”) selectively recognizes target not occupied by the inhibitor, and a second pair recognizes both occupied and unoccupied target (“TOTAL-RIPK1”). A drop in the normalized FREE-RIPK1 detection signal therefore is a relative measure of occupancy in a sample such as blood. As the TEAR1 assay relies on competition between the inhibitor and an antibody pair, and does not involve sample heating, it is an orthogonal measure of RIPK1 occupancy to the CETSA® assays.

Testing the same compounds, we found a strong correlation between the EC_50_ values for the two assays, with a Pearson correlation of 0.82 (Fig. 4B, S8C, and Table 1). Interestingly, the original report of the TEAR1 assay mentioned that necrostatin-type inhibitors showed no response, although we found three necrostatins (1, 1S, and 4) but not necrostatin 5 had measurable TEAR1 EC_50_ values in two biological replicates in our hands.

All together, these evaluations and data demonstrate that CETSA® can be a robust and sensitive assay for quantifying target engagement from small volumes of human whole blood.

## Discussion

We have demonstrated that Alpha CETSA® and MSD CETSA® show excellent potential for quantitative CETSA® engagement preclinically or clinically in whole blood. In our optimized protocols, both assays are sensitive, robust, and can be paused after lysis. Importantly, each protocol requires less than 100 μL of blood per well.

In the case of our proof-of-concept target RIPK1, the inter-donor *T*_*m*_ range is small relative to the large thermal stabilization induced by inhibitors such as Takeda compound 22. This large stabilization clearly contributes to the high S/B and Z’ values we are able to measure with Alpha CETSA® and MSD CETSA®. Whether any other targets besides RIPK1 are suitable for CETSA® will require similar validation. Some targets and target-compound pairs have more modest stabilization or destabilization effects that may be suitable for preclinical mechanism of action validation, but not for robust and quantitative preclinical or clinical assays. As an illustrative example, CETSA®-MS studies with the promiscuous inhibitor staurosporine have shown variable stabilization or destabilization across kinase targets. Although the CETSA® EC_50_ values correlate with staurosporine affinities (25), the effect size is more target-dependent and in some cases, highly occupied targets can have small or no CETSA® effects (26, 27).

Another factor that may influence the suitability of quantitative CETSA® in whole blood is the target’s intrinsic sensitivity to differences in biological state. Previous studies have shown measurable CETSA® effects for some targets due to changes including post-translational modifications, protein-protein interactions, and substrate metabolite levels (reviewed by ref. 28). While RIPK1 has no frequent protein-coding variants in healthy populations (29), we observed a difference in the Δ*T*_*m*_ for compound 22 in whole blood compared to isolated PBMCs. Although the underlying reason is unclear, possible explanations include a non-PBMC immune cell population in whole blood, such as neutrophils, with a different state of RIPK1, or with greater uptake of the compound.

We also found that the antibody pair selected for a plate-based CETSA® assay can influence the readout via a compound-dependent reduction in the target quantification in the absence of sample heating. Although the source of this non-CETSA® effect is difficult to determine, in some cases it may be due to compound-dependent conformational changes altering antibody binding of the target epitope – one extreme example of this is the MSD-based TEAR1 assay, where a complete loss of epitope detection is used to infer target engagement. For antibody pairs where the signal is not absolutely competed or reduced by the compound, including unheated samples with compound treatment can be used to correct for this effect. Ideally, using an antibody pair without such a non-CETSA® effect is recommended.

While multiple assays exist for measuring direct or proximal target engagement, CETSA® offers some unique characteristics that make it a useful complement to current approaches. Most established assays for whole blood focus on quantifying the occupancy of covalent inhibitors or reversible inhibitors with slow off-rates that are less sensitive to cell lysis steps (30). The key “heat shock” step of CETSA® is performed on intact cells or tissue, thus decreasing sensitivity to the kinetics of a binding interaction relative to current approaches. This is particularly important for the accurate quantification of direct target engagement after compound dosing.

Additionally, as CETSA® simply relies on the detection of total (soluble) target in a sample after heating, there are more reagents potentially available to detect a given target compared to one of its specific phosphorylation sites or other post-translational modifications, often the focus of proximal target engagement assays. We emphasize, however, that direct and proximal engagement assays are complementary and ideally both can be developed to evaluate a therapeutic hypothesis at preclinical and early clinical stages.

Finally, whole blood CETSA® can be implemented in preclinical animal models, including dogs and non-human primates. Although outside the scope of this study, we anticipate CETSA® could be used to determine dose-dependent target engagement in preclinical toxicology studies in comparison to compound pharmacokinetics and any dose-dependent adverse findings. Similarly, CETSA® could be useful, along with proximal target engagement assays, during dose optimization in early clinical development which has been a focus of recent regulatory initiatives such as the United States Food and Drug Administration’s Project Optimus (31,32).

The Alpha CETSA® and MSD CETSA® assays described here can be used for direct target engagement evaluation in whole blood. These assays can serve as an important roadmap for the implementation of whole blood CETSA® in various preclinical and clinical settings and provide an important tool for drug development.

## Acknowledgements

We would like to acknowledge assistance from the following AbbVie scientists: Dawn Bennett, Christian Goess, Michael Hoemann, and Xi Shi for helpful discussions about RIPK1 biology, reagents, and inhibitors; David Kinsman for synthesis of necrostatin-4; Paul Richardson, Heather Davis, and Haoyuan Zhang for assistance characterizing compound purity.

