## Supplementary Information for "Quantitative target engagement of RIPK1 in human whole blood via the cellular thermal shift assay for potential pre-clinical and clinical applications"

**Supplementary Methods**

**Cell Culture**

Jurkat cells were cultured in RPMI-1640 with 2 mM L-glutamine and supplemented with 10 mM HEPES, 1 mM sodium pyruvate, 4500 mg/L glucose, 1500 mg/L sodium bicarbonate, 1% penicillin-streptamycin, and 10% fetal bovine serum (FBS). RIPK1 knockout THP-1 cells (Abcam ab276121) and wildtype THP-1 cells (Abcam ab275477) were cultured in RPMI-1640 with 2 mM L-glutamine and supplemented with 1% penicillin-streptomycin, 10% FBS, and 0.05 mM 2-mercaptoethanol (Gibco 21985-023). PBMCs were cultured in RPMI-1640, supplemented with 1 % penicillin-streptomycin, 10% FBS (heat inactivated), and 1 µg/mL 2-mercaptoethanol (Gibco 21985-023). Cells were cultured in a humidified incubator at 37 °C, 5% $CO_2$ and verified to be free of mycoplasma contamination.

**Simple Western capillary immunoassay-based CETSA® with Jurkats**

Jurkat cells (60,000 cells per condition) were incubated with or without Takeda compound 22 at 37 °C for one hour followed by a 3 min CETSA heat shock at temperatures as indicated in Fig. 1B. Following lysis with NP-40 lysis buffer, samples were centrifuged at 20,000xg for 20 min and the soluble fraction isolated. Samples were then prepared for analysis on a Simple Western Wes capillary electrophoresis immunoassay system using a 12-230 kDa Separation Module (Bio-Techne SM-W004), following manufacturer protocols.

**Western blot-based CETSA® with human whole blood**

Fresh human heparinized blood was treated with vehicle or 10 µM Takeda compound 22, distributed into a MicroAmp Optical 96 well plate, and incubated for 1 h at 37 °C (total 3 mL of blood per condition in multiple wells). The appropriate wells were heat shocked for 3 min with temperature ranging from 37 to 61 °C on a Veriti thermal cycler. Blood samples were re-combined and PBMCs were isolated using the EasySep Direct Human PBMC Isolation Kit (STEMCELL, 19654) following manufacturer instructions. PBMC viability was determined using Trypan blue staining on Countess II FL automated cell counter (Thermo Fisher Scientific) and then lysed with NP-40 lysis buffer (8% NP-40 supplemented with 1KU benzonase, 1.5mM MgCl2 and Complete EDTA-free protease Inhibitor tablet, used as a 10X stock) by incubating on ice for 1 hour.

Samples were centrifuged at 20,000xg for 20 min and the soluble fraction was isolated and denatured at 95°C for 10min. Samples were run on NuPage 4–12% Bis-Tris Gels followed by transferred to a PVDF membrane using the iBlot2 system and iBlot gel PVDF transfer stacks. Membranes were incubated with a 1:1 mix of PBS and LiCor Intercept PBS blocking buffer for 1 hour at room temperature and overnight at 4°C with the primary antibody (1:1000 RIPK1, Abcam ab125072; or 1:1000 SOD1, Sigma-Aldrich HPA001401, as thermostable loading control). Blots were washed three times in PBS containing 0.1% Tween 20, followed by incubation with LI-COR secondary antibody at a 1:10000 dilution and imaged on the LICOR Odyssey CLx.

**MSD assay optimization**

MSD multiarray 96-well plates were coated overnight at 4°C with mouse anti-human RIPK1 capture antibody (BD Bioscience 610459) at the indicated concentrations in Fig. S1, as otherwise described in the Methods. Rabbit anti-human RIPK1 antibody (Abcam ab125072) was used for primary detection. Concentrations of each component of the antibody pair was optimized using Jurkat cells (60,000 cells per condition) heated at either 37 °C or 60 °C, testing for the highest electrochemiluminescence signal and signal/background ratio. Three different cell lysis buffers were also tested for optimal signal: Cell Signaling Technologies (CST) Cell Lysis Buffer (Cell Signaling Technologies 9803S) supplemented with Protease Inhibitor Cocktail (Millipore Sigma P8340), MSD Tris Lysis Buffer (Meso Scale Discovery R60TX-2), and NP-40 lysis buffer (8% NP-40 supplemented with 1KU benzonase, 1.5mM $MgCl_2$ and Complete EDTA-free protease Inhibitor tablet). All lysis buffers were used as 10X stock solutions.

To determine MSD signal linearity, whole blood was heated at 61 °C for 3 minutes to ensure complete denaturation of native RIPK1. This blood was then used as a medium for titration of recombinant RIPK1 (Abnova H00008737-P01) ranging from 0.1 pM to 10nM.

**MSD and Alpha CETSA® with cells**

Isolated PBMCs, THP-1 wildtype, or THP-1 RIPK1 knockout cells (at indicated cell numbers for PBMCs or 60,000 cells for THP-1s) were incubated with or without Takeda compound 22 at 37 °C for one hour followed by a 3 min CETSA® heat shock – at temperatures for PBMCs in Fig. 1, or at 3 7°C or 61 °C for THP-1s. Cells were then lysed with CETSA® lysis buffer 3 (for Alpha CETSA®) or NP-40 lysis buffer (for MSD CETSA®). Subsequent protocol steps for MSD and Alpha CETSA® were performed as described in the methods.

### Supplementary Figures

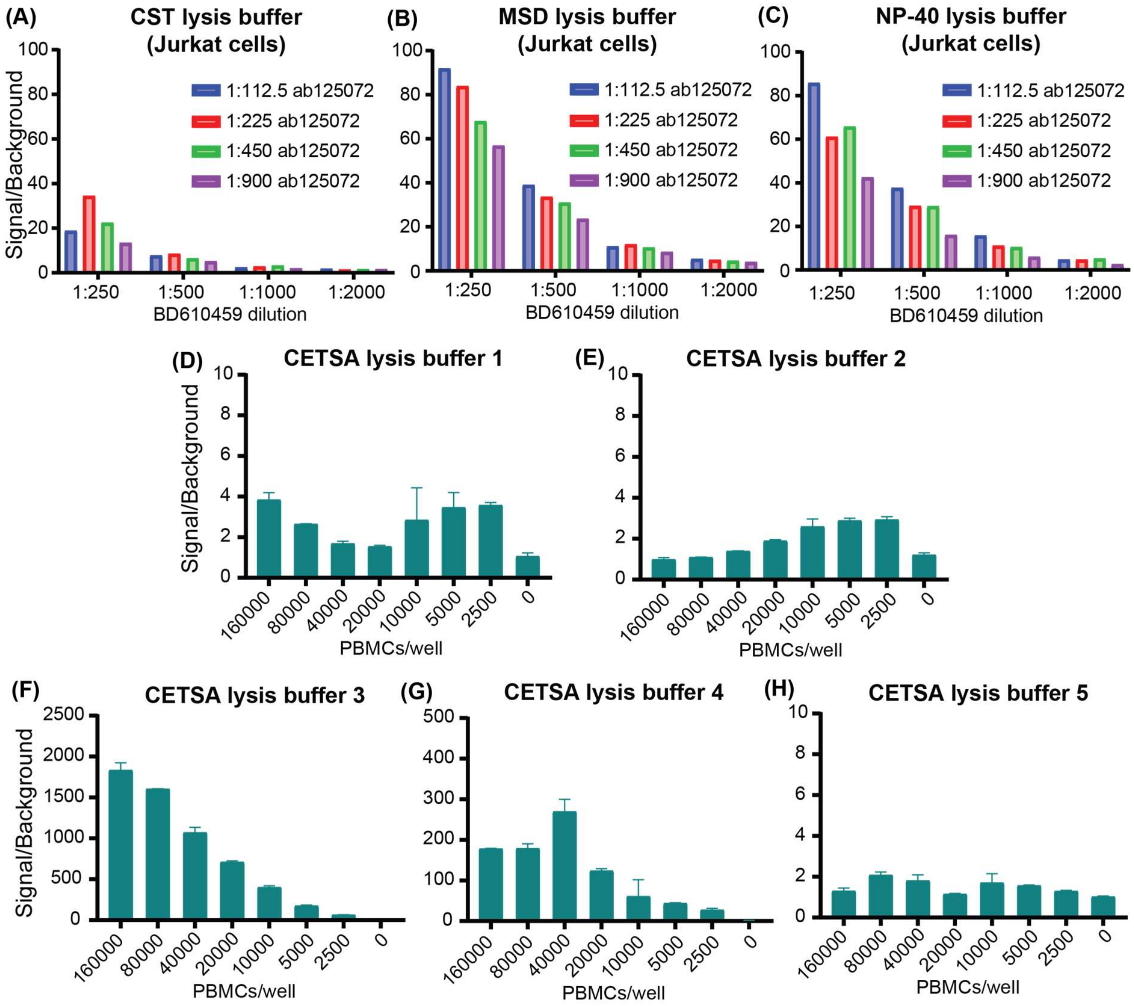


**Supplementary Figure 1.** Optimization of Alpha and MSD CETSA® assays. (A-C) RIPK1 MSD CETSA® assay signal-to-background (S/B) ratio for Jurkat cells with different lysis buffers (CST, MSD Tris, or NP-40 lysis buffers, see SI Methods), BD610459 capture antibody concentrations, and ab125072 detection antibody concentrations. (D-H) RIPK1 Alpha CETSA® was optimized using isolated PBMCs with Perkin Elmer CETSA® cell lysis buffers 1 to 5, revealing maximum signal with Buffer 3.

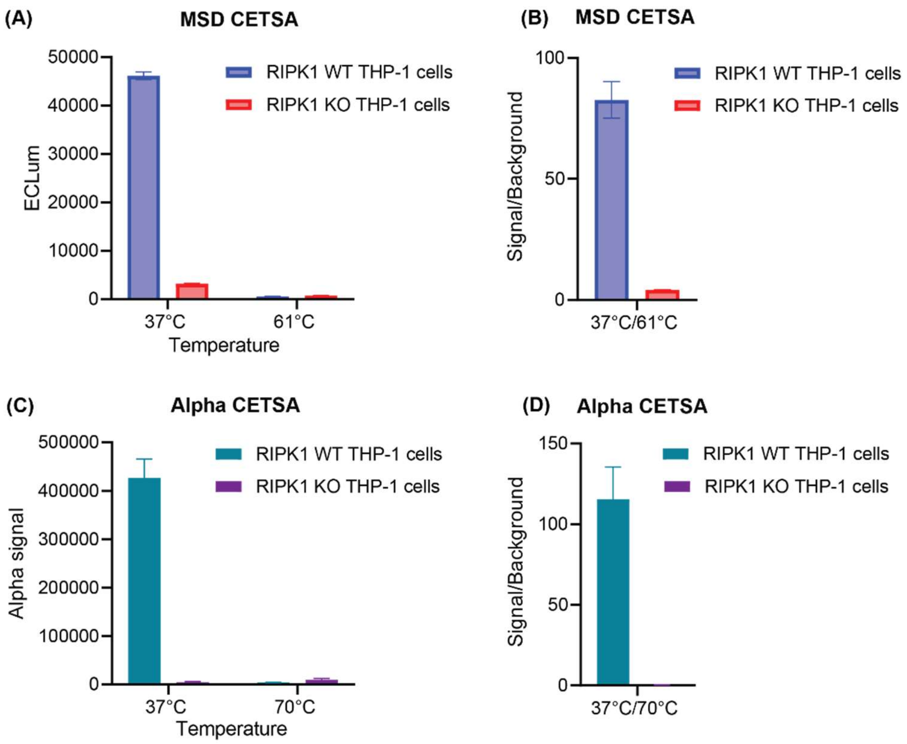


**Supplementary Figure 2.** Validation of assay specificity for RIPK1. MSD CETSA® (A,B) and Alpha CETSA® assay data (C,D) with isogenic RIPK1 wildtype (WT) and knockout (KO) THP-1 cells show that, for each assay, signal is specific to RIPK1 and can be abrogated with heating at 61 °C. Data represents mean ± standard error (SE), N=6 for MSD CETSA® and N=8 for Alpha CETSA®.

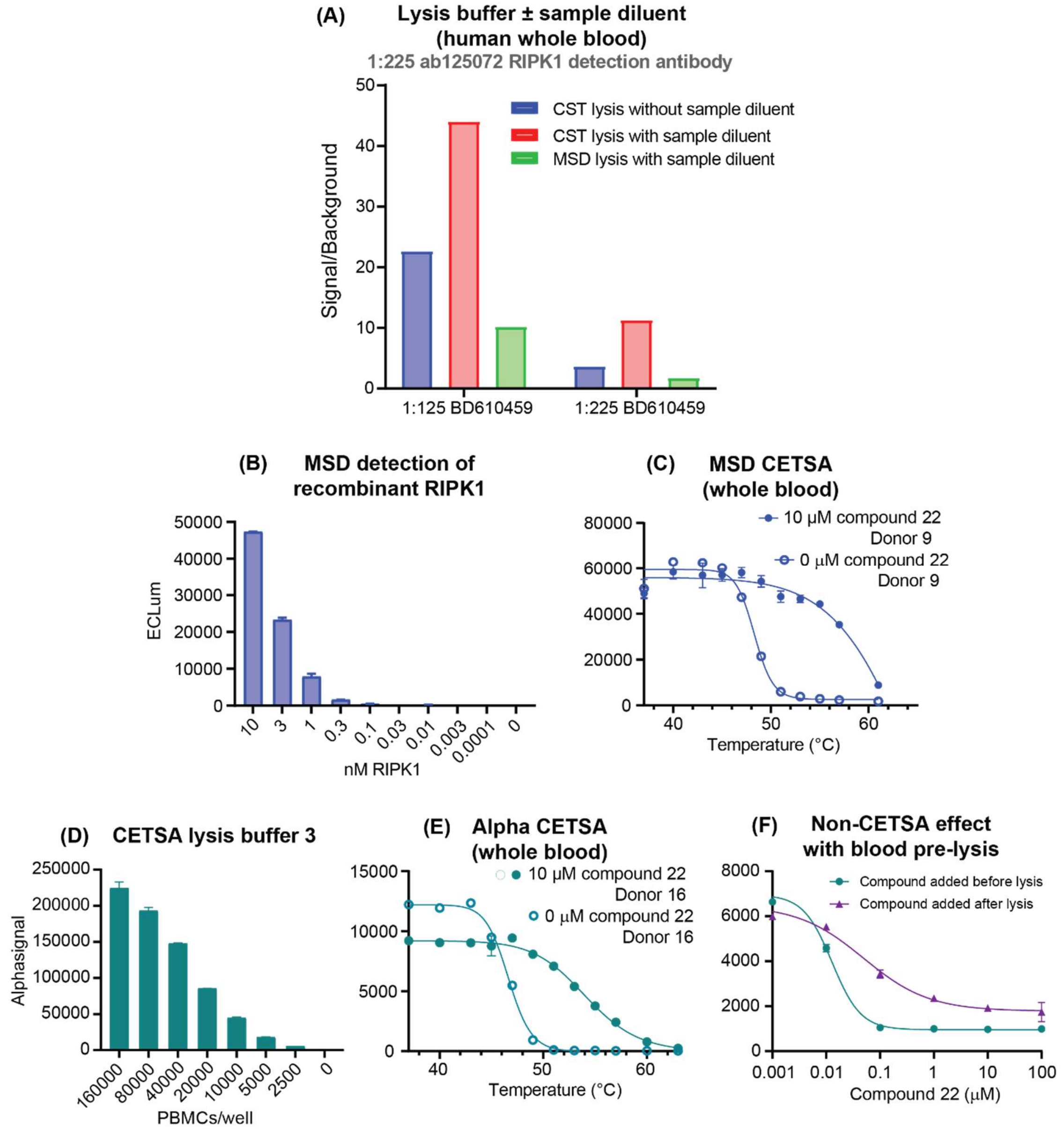


**Supplementary Figure 3.** Assay optimization and performance in whole blood. (A) MSD CETSA® S/B for human whole blood is improved by using CST lysis buffer, in contrast to experiments with Jurkat cells (Fig. S1). Diluting blood 1:1 with sample diluent further increases S/B. (B) MSD CETSA® signal for a dilution series of RIPK1 recombinant protein spiked into blood with fully denatured endogenous RIPK1, showing the assay is within its linear range at a signal intensity for a representative experiment with human whole blood (C, raw data for Fig. 1E). Note the minimal reduction in RIPK1 detection signal in the absence of heating (37 °C) and the presence of Takeda compound 22 (10 μM) – the antibody pair's "non-CETSA® effect". (D) Alpha CETSA® signal with a varying number of PBMCs shows the assay is within its linear range at a signal intensity for a representative experiment with human whole blood (E, raw data for Fig. 1F). Note the moderate reduction of RIPK1 detection signal (non-CETSA® effect) in the absence of heating and the presence of Takeda compound 22. (F) A non-CETSA® effect is observed for Alpha CETSA® when compound is added after blood lysis, suggesting it is not due to a compound-dependent reduction in RIPK1 abundance. Error bars for (C) and (E) were derived from technical replicates.

| Variability | Reading | Average | SE | %CV | n |
|---|---|---|---|---|---|
| **S/B Alpha CETSA® (at Pelago)** | Alphasignal 37°C/70°C | 272.44 | 78.20 | 28.70 | n=6 |
| **S/B Alpha CETSA® (at AbbVie)** | Alphasignal 37°C/70°C | 154.99 | 21.54 | 13.90 | n=10 |
| **S/B MSD CETSA® (at AbbVie)** | ECLum 37°C/61°C | 60.12 | 9.56 | 15.90 | n=33 |
| **Z' Alpha CETSA® (at Pelago)** | Z' | 0.76 | 0.05 | 6.74 | n=6 |
| **Z' Alpha CETSA® (at AbbVie)** | Z' | 0.59 | 0.10 | 16.12 | n=10 |
| **Z' MSD CETSA® (at AbbVie)** | Z' | 0.78 | 0.02 | 2.98 | n=33 |
| **MSD inter-donor and inter-run variability** | ECLum 37°C | 45999.46 | 1228.68 | 2.67 | n=135 |
| | ECLum 61°C | 1285.42 | 95.33 | 7.42 | n=113 |
| **MSD intra-donor and inter-run variability** | ECLum 37°C | 37347.03 | 2221.76 | 5.91 | n=43 |
| | ECLum 61°C | 686.48 | 40.72 | 7.29 | n=38 |
| **MSD intra-donor and day variability** | ECLum 37°C | 42600.92 | 1913.30 | 4.70 | n=80 |
| | ECLum 61°C | 1195.76 | 27.04 | 2.86 | n=74 |

**Supplemental Table 1.** Average signal-to-background (S/B) ratios and Z' values for the MSD and Alpha CETSA® assays across multiple runs or wells (n). "Signal" (S) corresponds to unheated (37 °C) blood sample replicates and "background" (B) to blood sample replicates heated at the maximal temperature (61 °C for MSD, 70 °C for Alpha). ECLum refers to the MSD assay signal. Due to a consistent use of anonymized donor codes at AbbVie, the average, standard error (SE), and coefficient of variation (CV) signal and background values could be determined for the MSD CETSA® assay across: different donors on different runs (inter-donor and inter-run), the same donors on different plates/runs (intra-donor and inter-run), and the same donors on different days or blood draws (intra-donor and inter-day).

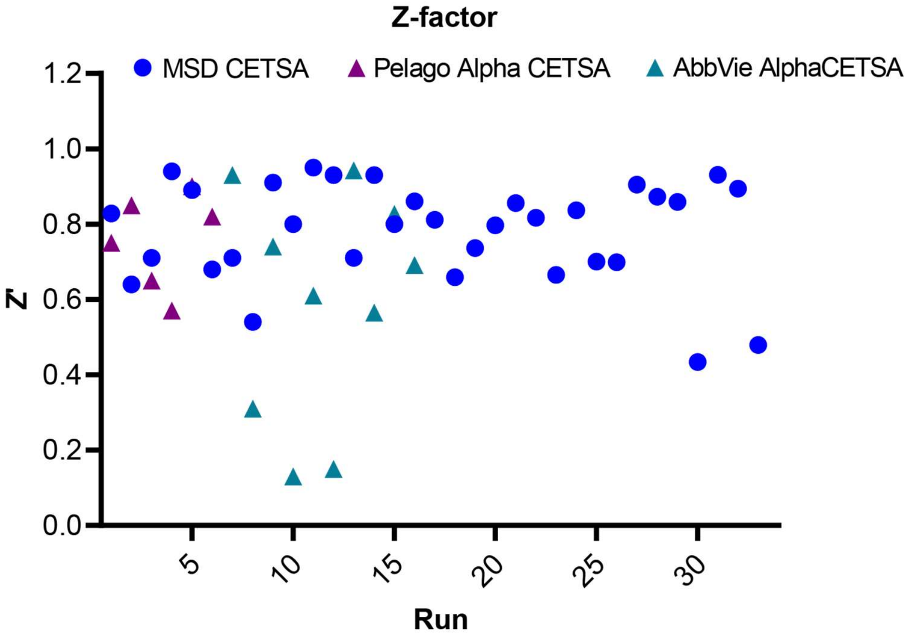


**Supplementary Figure 4.** MSD and Alpha CETSA® Z' measurements with human whole blood, determined for each run (plate) from signal for unheated (37 °C) blood sample replicates and for blood sample replicates heated at the maximal temperature (61 °C for MSD, 70 °C for Alpha). Alpha CETSA® Z' measurements are separated into assay runs at Pelago (cyan triangles) and AbbVie (magenta triangles), while all MSD CETSA® (blue circles) were performed at AbbVie.

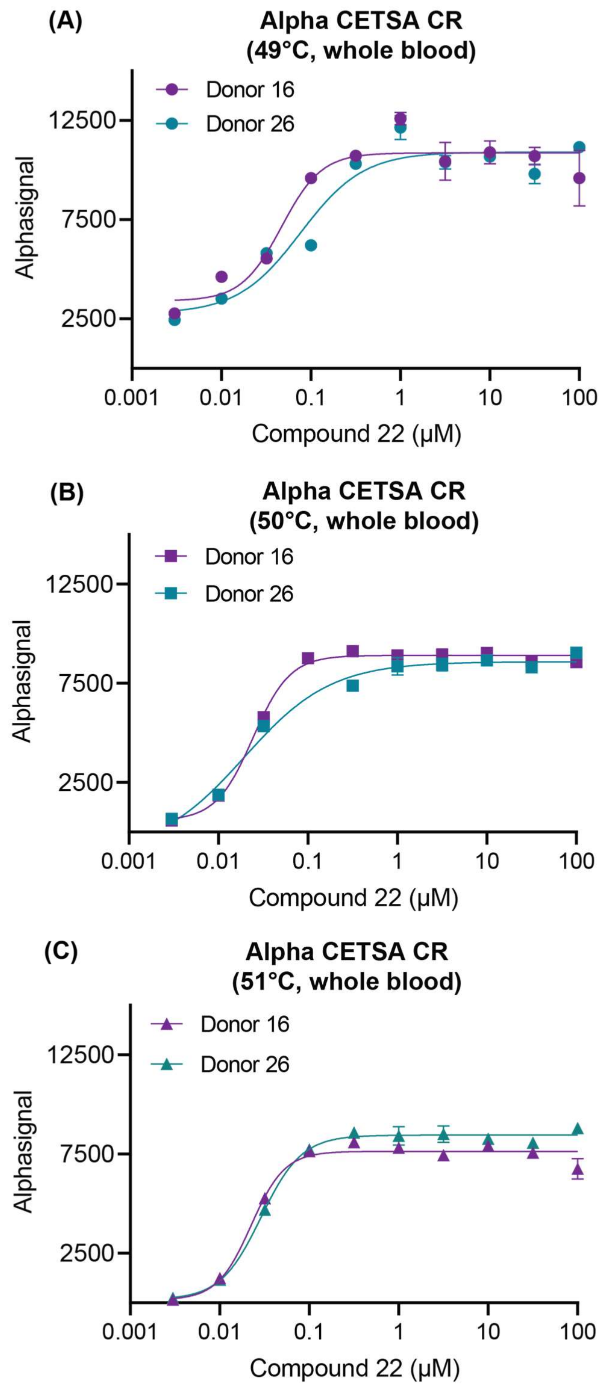


**Supplementary Figure 5.** Concentration-response (CR) Alpha CETSA® experiments with Takeda compound 22 show similar signal windows for two donors at three different temperatures, 49 °C, 50 °C, and 51 °C.

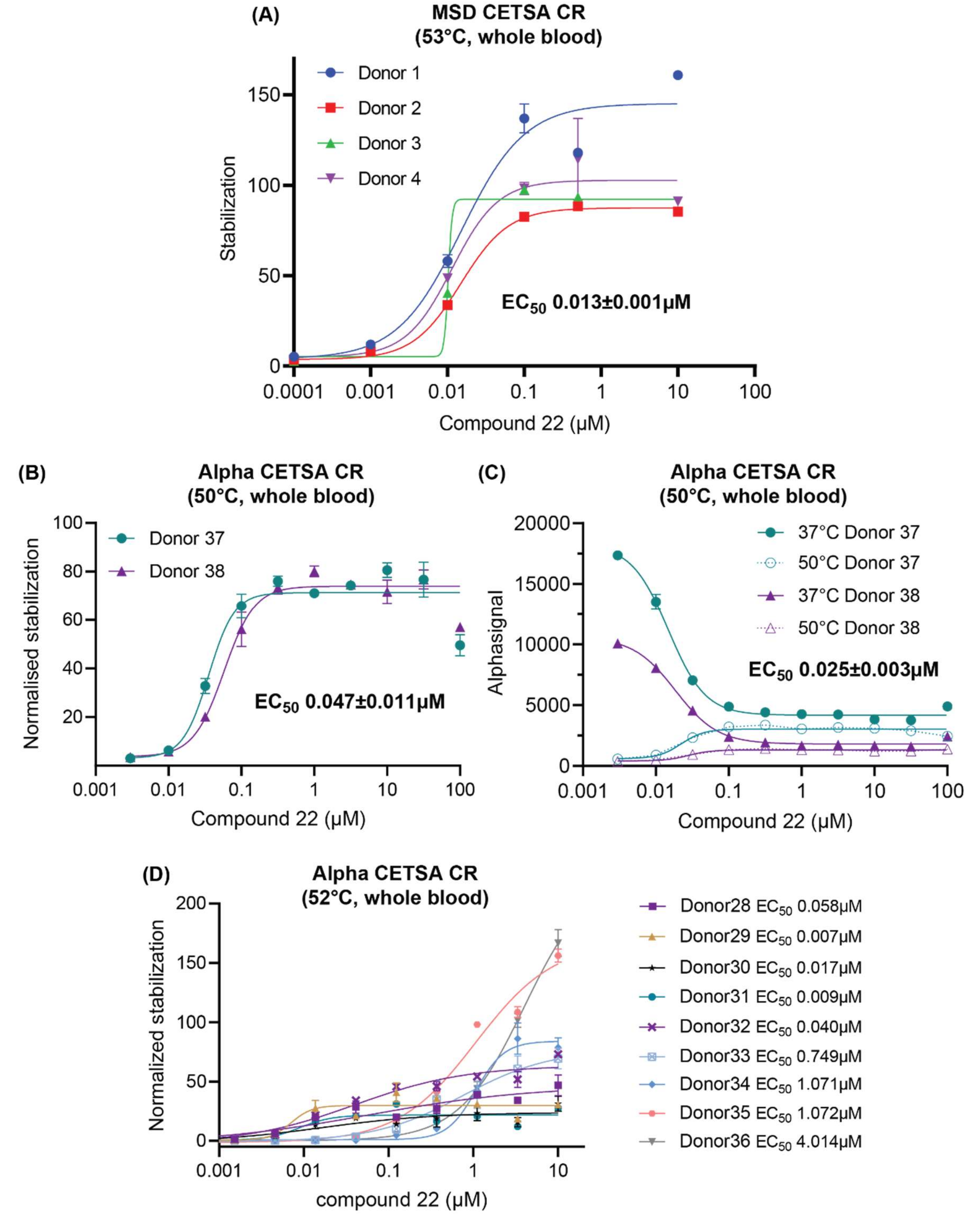


**Supplementary Figure 6.** Assay non-CETSA® effects in concentration-response (CR) experiments. (A) CR MSD CETSA® data for Takeda compound 22 without a correction for the compound-dependent non-CETSA® reduction in signal. There is minimal change in signal or calculated $EC_{50}$ (compare to corresponding corrected data from the same experiment in Fig. 3B). CR Alpha CETSA® data for compound 22 with two additional donors with (B) or without (C) a correction for the non-CETSA® effect reveals a significant compound-dependent reduction in RIPK1 signal in the absence of heating (37 °C curves in C). Despite this, the calculated $EC_{50}$ was only modestly affected. (D) CR Alpha CETSA® experiments with compound 22 at AbbVie show a non-CETSA® effect of similar magnitude to in C, but greater variability in $EC_{50}$ and Z' (SI Table 1).

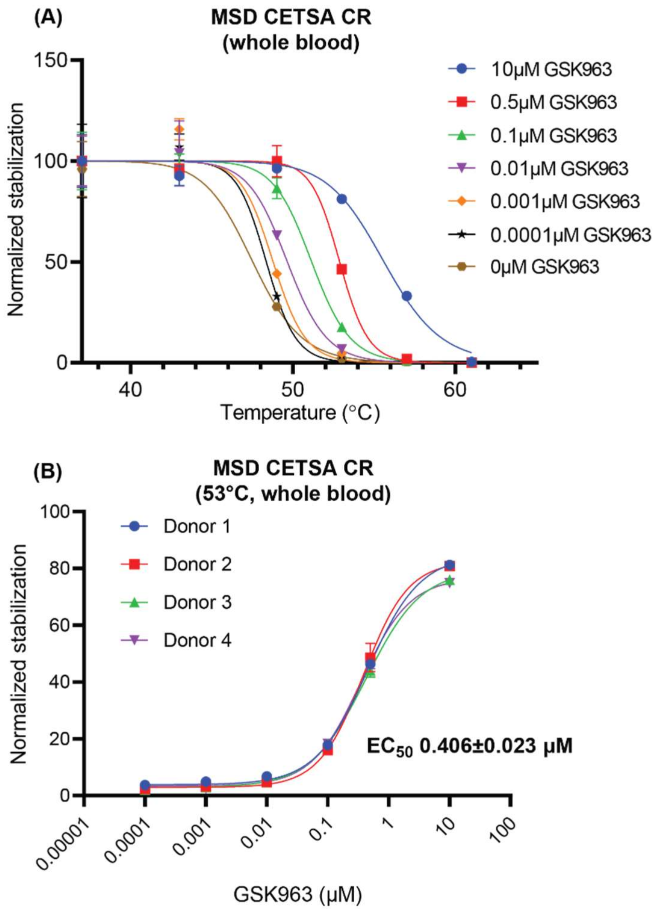


**Supplementary Figure 7.** Additional MSD CETSA® data for GSK'963. (A) Multiple concentration, multiple temperature data. (B) CR data with GSK'963 at 53 °C.

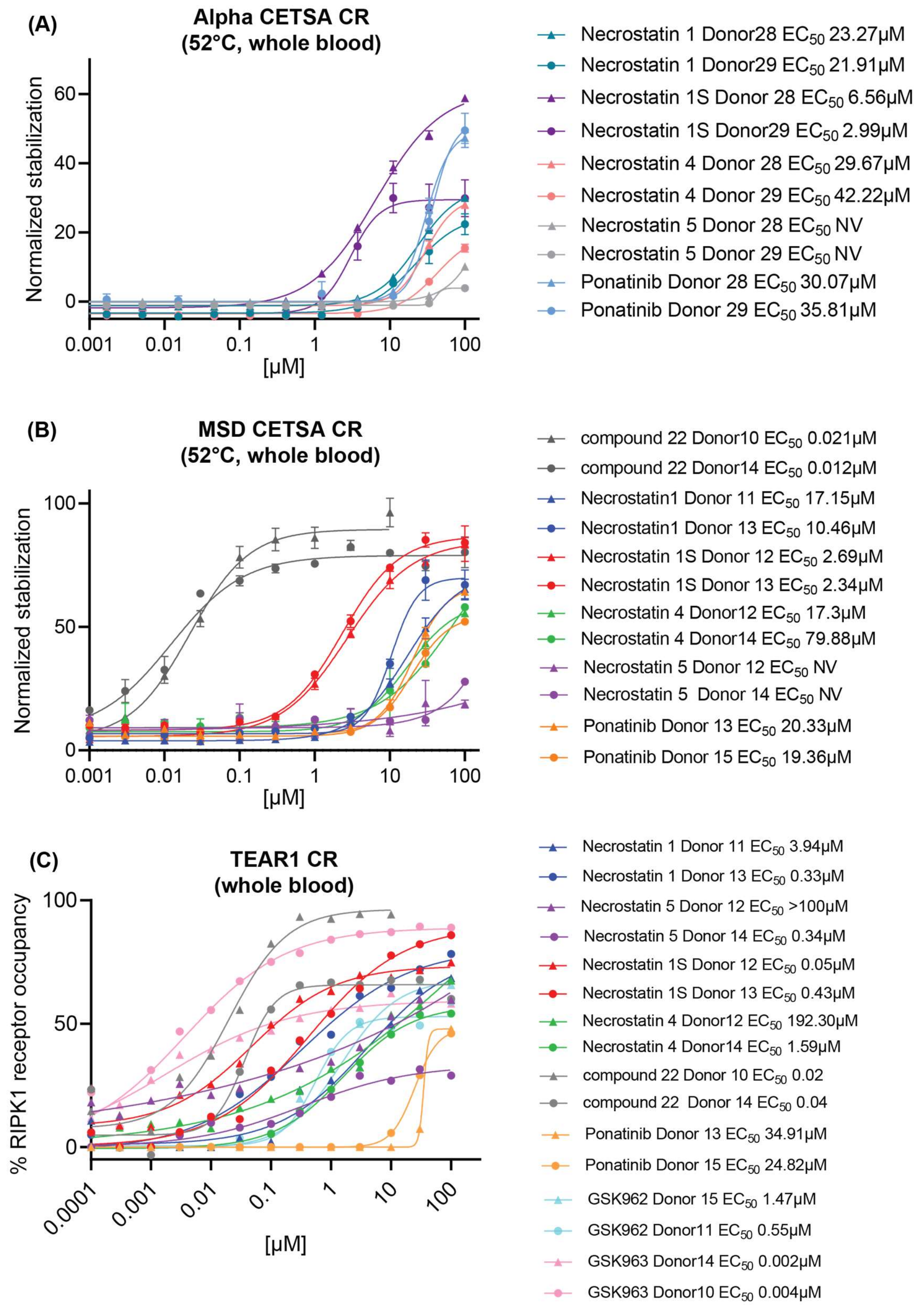


**Supplementary Figure 8.** Direct engagement data for additional compounds in whole blood: (A) Alpha CETSA®, (B) MSD CETSA®, and (C) TEAR1 assay.
